# Chimeric Antigen Cytotoxic Receptors for In-Vivo Engineering of Tumor-targeting Natural Killer Cells

**DOI:** 10.1101/2023.11.07.565144

**Authors:** Neha Diwanji, Daniel Getts, Yuxiao Wang

**Affiliations:** Myeloid Therapeutics, Cambridge, MA

## Abstract

Ex vivo chimeric antigen receptor (CAR) NK cells face challenges in manufacturing, and have limited tumor infiltration and in vivo persistency. A method leveraging mRNA-based delivery for in-vivo engineering of human NK cells could address these issues but has not been established. Here we developed an in-vivo NK cell engineering method by designing CARs that capitalize on inherent NK receptor biology for specific expression and function. These CARs utilize the Immunoreceptor Tyrosine-based Activation Motif (ITAM)-containing signaling adaptor in human NK cells for tumor destruction and cytokine response. We demonstrated that an NKp44-based CAR’s expression and function depend on the signaling adaptor DAP12. This approach enables precise mRNA-driven in-vivo NK cell programming against tumors, ensuring specificity and reducing off-target expression in non-immune healthy tissues.

## INTRODUCTION

Natural killer (NK) cells are a subset of innate lymphoid cells that can recognize and eliminate infected, stressed, and malignant cells. They play important roles in innate immune surveillance of cancer, and can produce cytokines and chemokines to generate robust adaptive immune responses (1). Unlike T cells, NK cells activate in a human leukocyte antigen (HLA) independent manner and can initiate tumor killing without causing potential graft-versus-host disease. NK cells express a large variety of inhibitory and activating receptors. Interactions between these receptors and their ligands trigger signaling pathways that collectively determine the functional outcome between inhibition and activation (2, 3). Extensive translational research and clinical development have focused on ex vivo engineering of NK cells with a chimeric antigen receptor (CAR) that recognizes tumor antigens to override inhibitory signals and initiates antigen specific tumor killing. This approach has shown preliminary clinical efficacy in hematological tumors but is faced with numerous challenges and limitations (4). NK cells are resistant to viral transduction, difficult to expand ex vivo and to cryopreserve, all of which increase costs and the complexity of product development and delivery (5, 6). Finally, ex vivo expanded NK cells suffer from short persistency in patients and low efficiency in tumor infiltration, as well as susceptibility to the immune suppressive tumor microenvironment (TME) (6). The ability to program NK cells in situ, without the need for ex-vivo manipulation may overcome these challenges, allowing their full capability to be harnessed for cancer treatment.

Messenger RNA (mRNA) therapeutics are rapidly growing thanks to the rapid development and approval of COVID-19 mRNA vaccines (7, 8). Advancements in mRNA sequence optimization, synthesis and delivery methods provide opportunities for direct engineering and programming of immune cells in vivo (9). Compared with current ex vivo cell therapy approaches, mRNA-based in vivo engineering approach has numerous advantages including simple, low-cost manufacturing, avoidance of insertional mutagenesis that are associated with viral integration, and flexibility in dosing and combining with other modalities such as checkpoint blockade. Conventional mRNA delivery approaches via lipid nanoparticles (LNP) when infused systemically result in significant accumulation in liver tissue (10). Expression of a functional CAR in liver could contribute to toxicity due to tonic proinflammatory signaling, therefore conventional mRNA/LNP has only been used for ex vivo transfection of immune cells (11). Although lipid chemistry optimization reportedly can promote LNP uptake by immune cells, the safety profiles of these novel lipids remain unclear and with only marginal gains in delivery specificity (12, 13).

Several NK activating receptors including NKG2D and the Natural Cytotoxicity Receptor (NCR) family that includes NKp30, NKp44 and NKp46 have been repurposed for the design of CARs for NK cells and T cells (14). However, the current generation of NK activating receptor-based CARs largely resembles the design of CAR-T constructs. Generally, the design involves fusion of ITAM-containing signaling adaptor domain from CD3ζ or DAP12, with a co-stimulatory domain from 41BB or CD28, while retaining the native extracellular domains (ECD) that recognize natural ligands (14). These CAR constructs were used to produce CAR-T or CAR-NK cells ex vivo and showed tumor cell killing activities. Nevertheless, delivery of such constructs in vivo using conventional mRNA/LNP will result in CAR expression in all different cell types that internalize LNP and translate the CAR mRNA, posing specificity issues and toxicity risks.

In this study we present a novel approach that harnesses the natural expression and functional properties of NK activating receptors to achieve specific expression of a tumor-directed CAR in NK cells and overcome the above-mentioned limitations. Many NK activating receptors lack intracellular signaling motifs, and instead form heterodimers with a signaling adaptor such as Fc receptor Y-chain (FcRY), CD3ζ and DAP12 for activation and downstream signaling. Formation of the receptor complex is mediated by interactions between a positively charged amino acid in the transmembrane domain of the NK activating receptors and a negatively charged residue in that of the signaling adaptor (15). Intriguingly, studies have shown that for some of the activating receptors, their cell surface expression also requires the presence of the signaling adaptor. For example, NK cells lacking the expression of FcRY showed greatly reduced expression of NKp30 and NKp46 (16, 17). In addition, NKp44 failed to express in Jurkat cells without co-transfection of DAP12 (18). The exact mechanism for such expression dependency remains unclear but could be due to reduced protein stability when the positively charged residue in the transmembrane domain is not compensated by interacting with the negatively charged residue in the signaling adaptor. We hypothesized that this could be harnessed to design a CAR that can only be stably expressed in NK cells but not in other cell types that lack the cognate signaling adaptor. This would enable the ability to systemically deliver the CAR via conventional mRNA/LNP and avoid expression in non-immune tissues that generally do not express signaling adaptors such as CD3ζ and DAP12. To achieve this, we designed and optimized a series of CARs based on the multiple activating receptors from NK cells. Armed with an scFv derived from the anti-HER2 antibody trastuzumab, these novel CAR constructs when expressed in NK cells can recognize HER2+ tumor and showed potent tumor killing activity as well as induction of cytokines and chemokines. Using the hepatocellular carcinoma line Huh7 as a model for hepatocytes, we determined if CAR expression requires their corresponding signaling adaptor and found that NKp44-based CAR depended on the adaptor DAP12 for its surface expression. Collectively, these results support further clinical development of NKp44-based CAR for in vivo programming of cancer-targeting NK cells.

## MATERIALS AND METHODS

### Cells

Huh7, Raji, and SKOV3 cells were purchased from the American Type Culture Collection. Huh7 cells were maintained in Dulbecco’s modified Eagle medium with GlutaMAX supplement (Gibco), 10% fetal bovine serum (FBS) (Gibco) and 1% penicillin–streptomycin–L-glutamine (PSG) (Gibco). Raji cells were maintained in suspension in Roswell Park Memorial Institute medium (Gibco) supplemented with 10% FBS and 1% PSG. SKOV3 cells were maintained in McCoy’s 5A media (Gibco) supplemented with 10% FBS and 1% PSG. EasySep™ Human NK cell-negative isolation kit (StemCell Technologies) was used to isolate human peripheral blood NK cells from healthy donor leukopaks. ImmunoCult™ NK cell-expansion kit (StemCell Technologies) was used following the manufacturer’s recommendations to activate isolated NK cells in vitro for 6 days. EasySep™ Human T-cell-negative isolation kit (StemCell Technologies) was used to isolate peripheral blood T cells from healthy donor leukopaks. ImmunoCult™ human CD3/CD28/CD2 T-cell activator (StemCell Technologies) was used to activate isolated T cells *in vitro* for 2 days following the manufacturer’s recommendations.

### Plasmid construction and in vitro transcription

Standard molecular biology techniques were used to clone all CAR constructs into the plasmid pcDNA3.1 backbone. All cloning steps were validated by restriction enzyme digestion and sequencing. GeneJet PCR purification kit (ThermoFisher Scientific) was used to linearize and purify the NK-CAR plasmids according to the manufacturer’s protocol. T7 RNA polymerase and CleanCap^®^ (TriLink) was used to transcribe mRNA from the linearized DNA template, poly(A) tail was added to the mRNA, and Monarch^®^ RNA purification kit (NEB) was used to purify the final mRNA according to the manufacturer’s protocol.

### Electroporation of NK cells or T cells

Activated NK cells or T cells from culture were resuspended in MaxCyte^®^ Electroporation buffer, and CAR mRNA (2 μg/mL) was added to the cells. The MaxCyte^®^ ATx system was used to electroporate the cells following the manufacturer’s recommended protocol. Following electroporation, the cells were recovered for 10 minutes in a 37°C cell culture incubator, after which the cells were added to prewarmed media and incubated at 37°C for 20 hours.

### Transfection of Huh7 cells

Huh7 cells were transfected with receptor mRNA using the Lipofectamine™ MessengerMAX™ (Invitrogen) mRNA transfection reagent. Transfection was performed with 100,000 cells, and NK-CAR mRNA prepared in Lipofectamine™ MessengerMAX™ according to the manufacturer’s protocols was added to the plated cells to obtain a final mRNA concentration of 0.5 μg/mL. The Lipofectamine™ mixture and mRNA were incubated with Huh7 cells for the indicated time, after which the cells were harvested and analyzed for surface expression of CAR. For co-transfection with signaling adaptors, 0.5 μg/mL of each mRNA was added to the cells for a total RNA concentration of 1 μg/mL. Specificity Index was calculated as the ratio of the average mean fluorescence intensity (MFI) of the CAR expression in the presence of a signaling adaptor to the average MFI of CAR expression in the absence of a signaling adaptor.

### Flow cytometry

A human HER2/ERBB2 protein-His tag (10004-H08H-100, Sino Biological) conjugated with Alexa Fluor 647 was used to determine the expression of HER2-CAR in transfected cells, and viability was determined by 7-AAD labeling. Anti-FLAG BV421 antibody with 7-AAD was used to determine the expression of CD19-CAR in transfected T cells. The antibodies used for surface phenotyping of NK cells and T cells included anti-CD56 PE-CY7 (5.1h11), anti-CD3 BV650 (OKT3), anti-CD4 BV785 (SK3), and anti-CD8a-APC-eFluor 780 (SK1). Cytek^®^ Northern Lights NL3000 flow cytometer was used to acquire cells, and FlowJo™ v10 (Becton Dickinson) was used to analyze the acquired data.

### In vitro cytotoxicity assay

SKOV3 tumor cells were used as targets in killing assay by control (Mock-transfected) or CAR- transfected NK cells at a 5:1 effector-to-target (E:T) ratio. The coculture was incubated at 37°C for 20 hours. CytoTox-Glo™ cytotoxicity assay kit (Promega) was used to measure the number of dead cells in the coculture population. Kit reagent was freshly prepared each time by adding AAF-Glo™ substrate to the assay buffer and was added to each well and incubated for 15 minutes at room temperature with orbital shaking. SpectraMax^®^ i3 (Molecular Devices) was used to measure luminescence. Effector-only wells, target-only wells, and total lysis of the target were included as technical controls. SKOV3 cytotoxicity was determined as follows: % Specific SKOV3 killing = [Average luminescence for NK+SKOV3 – Average luminescence for NK only – Average luminescence for SKOV3 only / Average luminescence for total lysis of SKOV3 – Average luminescence for SKOV3 only] × 100%. Raji tumor cells were used as targets in killing assay by control (Mock-transfected) or CAR-transfected T cells at a 5:1 E:T ratio. The Raji cells were labeled with CellTrace™ Violet and cocultured with T cells at 37°C for 40 hours. The coculture was stained with 7-AAD, and Raji-cell killing was calculated as the percentage increase in 7-AAD+ CellTrace™ Violet+ Raji cells.

### Luminex assay

NK-CAR and Mock NK cells were cocultured with SKOV3 tumor cells at a 5:1 E:T ratio for 24 hours at 37°C. Cytokine & Chemokine Convenience 34-Plex Human ProcartaPlex™ Panel 1A (Invitrogen) was used to collect and analyze the supernatant for levels of cytokines/chemokines following the protocol provided by the manufacturer. Data were acquired on a Luminex^®^200™, and xPonent^®^ software was used for analysis.

### Statistical analysis

Prism 9 software (GraphPad) was used to perform all statistical analyses. All central tendencies indicate the mean, and all error bars indicate the standard deviation (SD). Dunnett’s multiple comparisons test was used to generate ANOVA multiple-comparison *P* values. For all Fig.s, *P* values are indicated on the graphs.

## RESULTS AND DISCUSSION

### Design of CAR constructs based on NK activating receptors

Previous research showed that NK activating receptors including NKG2D, NKp30 and NKp46 can be fused with the CD3ζ intracellular signaling domain to express in T cells and mediate tumor killing (14). However, these CARs rely on the presence of NK receptor ligands on tumor surface for tumor recognition and activation, thus limiting their utility. We attempted to re-target their activities towards a specific tumor-associated antigen by fusing an extracellular tumor targeting scFv to the NK receptors, while preserving the native transmembrane and cytosolic domains for engagement of their signaling adaptors including FcRY, DAP10 and DAP12. To this end, activating receptors specific for NK cells were used to design 14 different CAR constructs (referred to as NK-CARs hereafter). The receptors chosen for the initial screening included the NCRs (19–22), and the NKG2 family of activating receptors NKG2C and NKG2D (23). All CARs were fused to an scFv derived from the anti-HER2 antibody Trastuzumab (24). Since the ECD of CAR tend to affect their expression and activity, the CARs were designed with ECD variations by taking the full-length, truncated ECD, or no ECD from the NK receptors (Fig. 1A-C). NK-CAR mRNAs were synthesized by *in vitro* transcription and transfected into human peripheral blood NK cells by electroporation. 20 hours after transfection, cell surface expression of the NK-CARs was examined by flow cytometry (Fig. 1A-C; Supplemental Fig 1). For NKp30-based CARs, HER2 scFv fused to full-length NKp30 showed the highest cell surface expression. Truncation or complete deletion of the ECD of NKp30 led to loss of cell surface expression of the CAR (Fig. 1A). NKp44-based CAR showed high cell surface expression for the full-length receptor as well as truncated ECD, however, the cell surface expression was lost when the ECD was deleted (Fig. 1B). The NKp46-based receptor demonstrated results similar to those of the NKp30-based CAR in which truncation or deletion of ECD caused loss of cell-surface expression of the CAR (Fig. 1C). These results indicate that the ECDs of the NCRs are important for maintaining cell surface expression of CARs. The NKG2C and NKG2D-based CARs did not show any cell surface expression for any format of the designed CARs (Supplemental Fig. 1). One reason for this could be due to that NKG2C and NKG2D are type II transmembrane proteins and fusion to scFv could pose challenges for their proper trafficking to the cell surface. However, inversion of the NKG2D domain orientation, to mimic that of a type I transmembrane protein (reverse NKG2D), also did not improve the cell surface expression of this CAR (Supplemental Fig. 1). Based on these results, we selected full-length versions of NKp30, NKp44 and NKp46 CAR designs for further studies.

**Figure 1:**
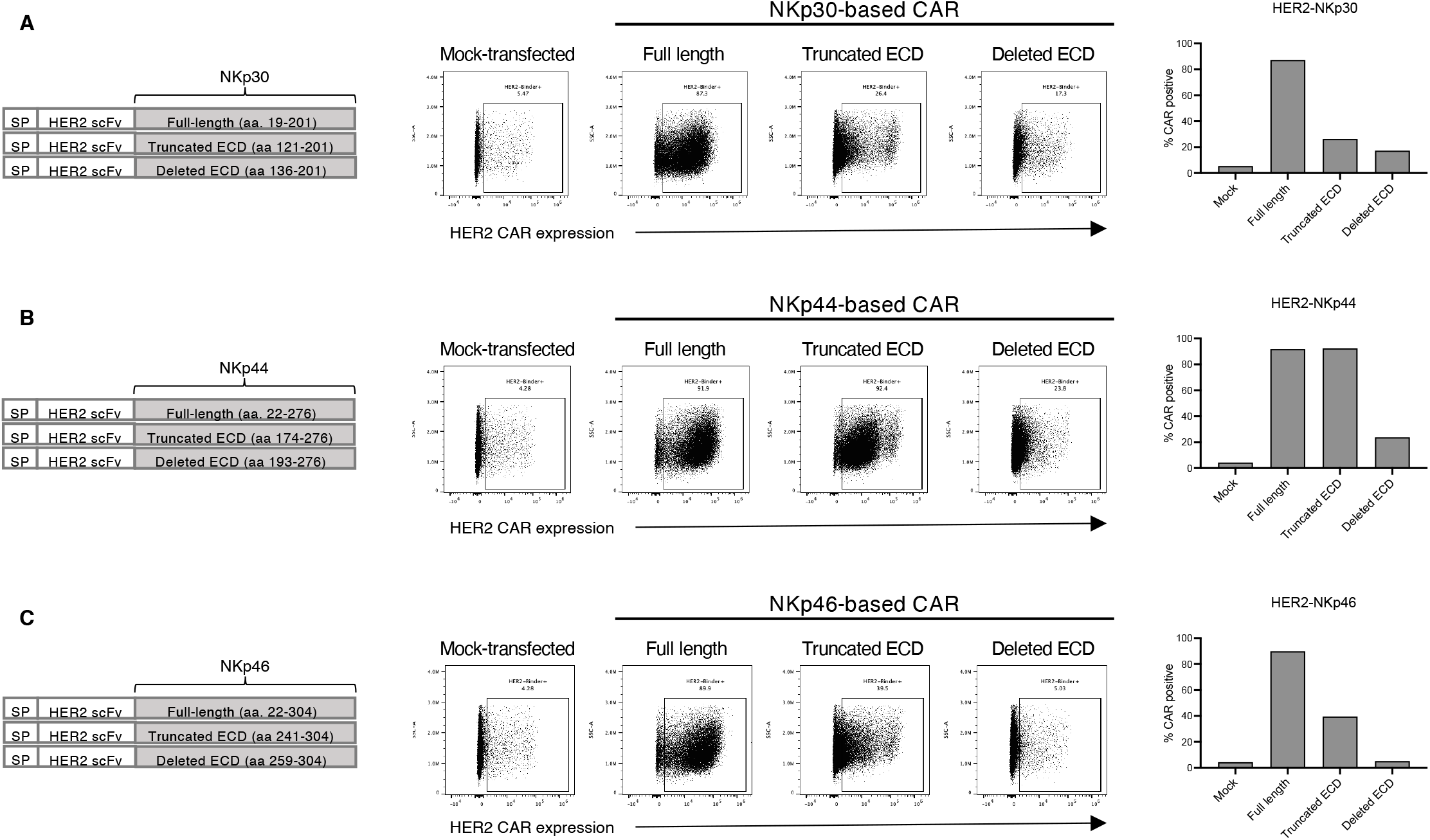
Design and expression of NK-CARs in human primary NK cells. Schematic representations of the designed NK-CARs and their cell-surface expressions in primary NK cells 20 hours after transfection are shown. (A) NKp30-based CAR; (B) NKp44-based CAR; (C) NKp46-based CAR. SP indicates signal peptide. Ranges of amino acids included in each constructs are shown. Data are representative of three independent experiments.

### Robust cell surface expression of NK-CARs relies on co-expression of corresponding signaling adaptors

NCRs associate with ITAM-containing signaling adaptors, such as CD3ζ, FcRγ and DAP12, for stable expression as well as downstream signaling after ligand stimulation. Therefore, we determined if cell-surface expression of NK-CARs depends on the presence of their cognate signaling adaptors, FcRγ for NKp30 and NKp46 and DAP12 for NKp44 (3). Identifying a receptor with expression conditional on the presence of an immune cell specific signaling adaptor would circumvent the problem of non-specific expression and activation of the CAR when delivered via mRNA/LNP in vivo. Since liver cells tend to take up LNP and express the mRNA encoded payload, we decided to use the hepatocellular carcinoma line Huh7 to examine the expression levels of the NK-CARs. Huh7 cells do not endogenously express FcRγ or DAP12 (25). We transfected Huh7 cells with NK-CAR mRNA with or without co-transfection with the cognate signaling adaptor mRNA. The result showed that cell surface expression of HER2- NKp30 CAR in Huh7 cells was high without FcRγ, and the presence of FcRγ only marginally increased the level of expression of HER2-NKp30 CAR (Fig. 2A). Similar results were observed for HER2-NKp46, although the level of expression increased substantially when FcRY was co- transfected (Fig. 2B). In contrast. HER2-NKp44 CAR strictly required the presence of cognate signaling subunit DAP12 for expression, and was not detectable on the cell surface in the absence of co-transfection with DAP12 (Fig. 2C). To quantify the result, the Specificity Index of expression was calculated by taking the ratio of mean fluorescence intensities (MFIs) of the receptors with or without co-transfection of the signaling adaptors. NKp30- and NKp46-based CARs showed modest levels of specificity, with up to a 3-fold increase in expression level when FcRY were co-transfected (Fig. 2D). In contrast, NKp44-based CAR showed a much higher specificity index (>10 fold) (Fig. 2D). These results together showed that among the NCRs, NKp44 has the highest level of dependency on the presence of cognate signaling adaptor for stable expression on the cell surface.

**Figure 2:**
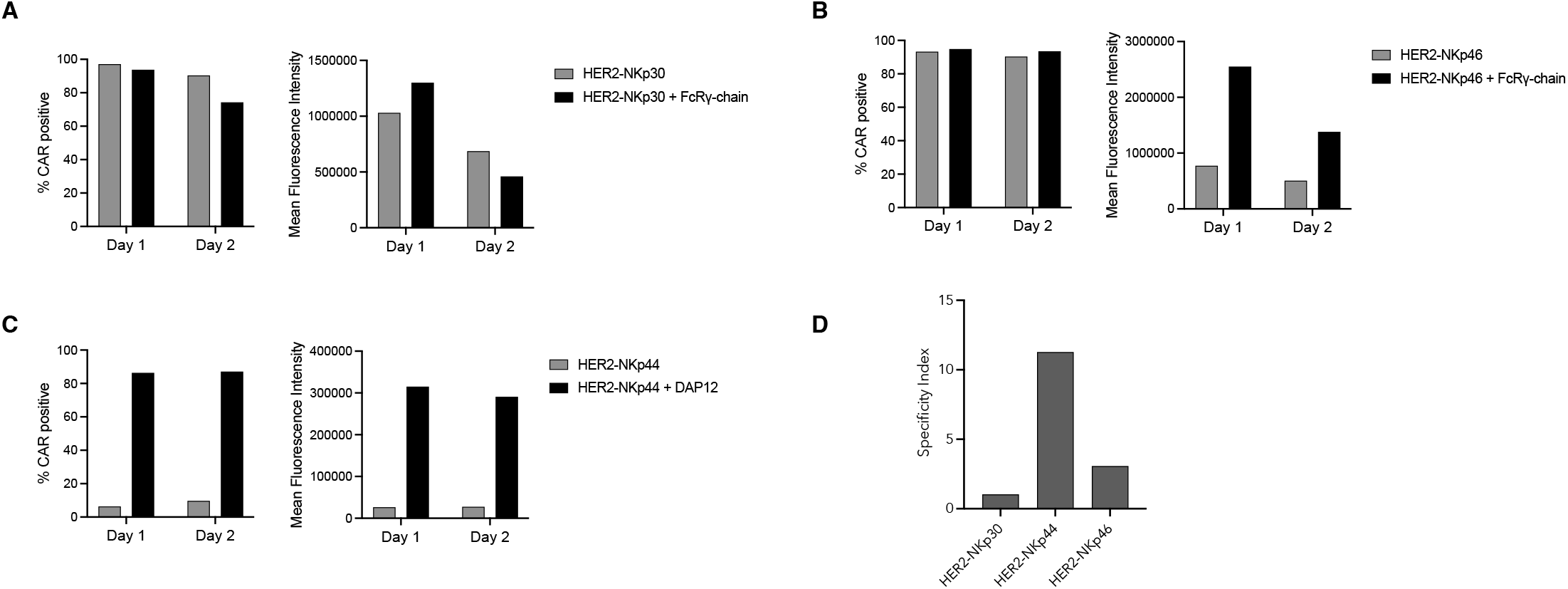
Cell surface expression levels of NK-CARs in the presence or absence of cognate signaling adaptors. NK-CAR mRNAs were transfected in Huh7 cells with and without co- transfection with the mRNA for signaling adaptors (A) HER2-NKp30 with and without FcRγ, (B) HER2-NKp46 with and without FcRγ and (C) HER2-NKp44 with and without DAP12. Each panel shows the percentage CAR positive and mean fluorescence intensity 1 or 2 days post transfection. (D) The Specificity Index of each CAR-NK constructs are calculated as the ratio of the surface expression of CAR in the presence of signaling adaptors to the surface expression in the absence of signaling adaptor. Representative data of two independent experiments are shown.

### Novel NK-CARs showed potent cytotoxic activity and cytokine response toward HER2+ cancer cells

We further evaluated the ability of NK-CARs to enhance NK-mediated killing of HER2+ tumor cells. We delivered the NK-CARs via mRNA to human primary NK cells, and cultured them with HER2+ SKOV3 ovarian cancer cells (Supplemental Fig. 2). Tumor cell death was measured using the CytoTox Glo assay (see Material and Method for details). To confirm that NK-CAR activity was antigen-specific and not due to existing NK ligands on tumor cell surface, we included mock-transfected NK cells as a negative control. For comparison, we also included a previously reported HER2 targeting CAR designed for NK cells, with CD3ζ as the intracellular activation signal and CD8a hinge and transmembrane domain (26). The cell surface expression of the CD3ζ-based CAR was comparable to the NCR-based CARs (Supplemental Fig. 3). All NK-CAR transfected cells displayed significant increase in SKOV3 cell lysis compared to mock-transfected NK cells. (Fig. 3A). Notably, the newly designed NK-CAR constructs performed similarly or slightly better than the CD3ζ based CAR. These results highlight the ability of NK- CARs to trigger natural activation pathway via engaging endogenous signaling adaptors, and promote targeted killing of tumor cells. To further evaluate the anti-tumor activity of NK-CARs, we measured the release of cytokines by the NK cells after co-culture with SKOV3 cells for 24 hours. All NK-CARs specifically enhanced expression of inflammatory cytokines TNF-alpha and IFN-gamma when co-cultured with SKOV3 (Fig. 3B). Additionally, chemokines, such as CCL2 and growth factor GM-CSF, were also significantly upregulated relative to mock-transfected NK cells by all of the NK-CAR constructs (Fig. 3B).

**Figure 3:**
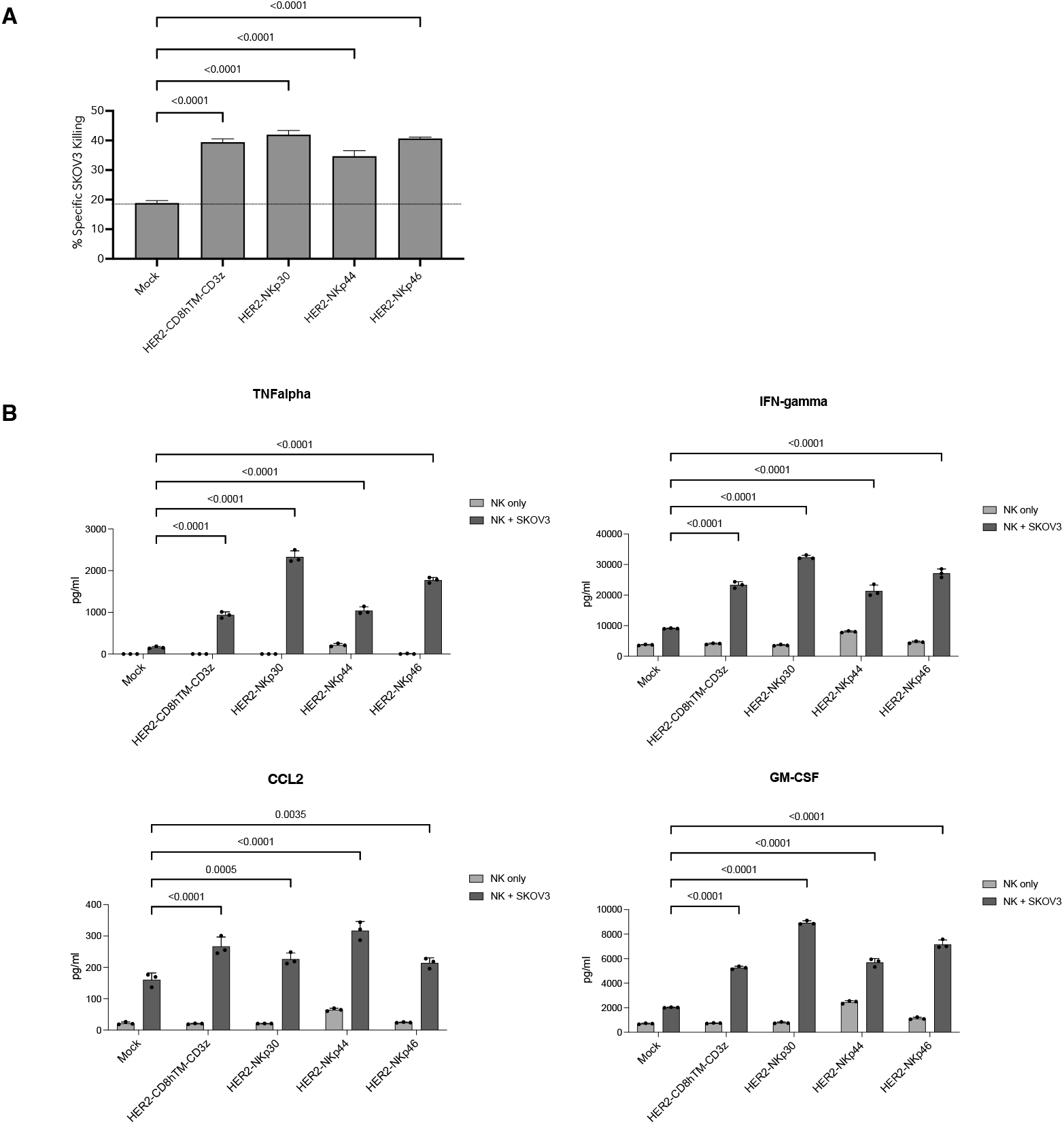
NK-CARs exhibit tumoricidal activity and elicit cytokine/chemokine response. (A) Peripheral blood NK cells transfected with HER2-targeting NK-CARs were cocultured with HER2+ SKOV3 tumor cells at a 5:1 effector-to-target (E:T) ratio for 20 hours at 37°C. The graph represents the percentage specific killing of SKOV3 cells. Data are representative of three independent experiments. The mean ± SD is plotted, and statistical significance is determined by ordinary one-way ANOVA with Dunnett’s multiple comparisons test between mock-transfected cells vs. NK-CAR transfected cells. *P* values are indicated on the graph. (B) TNF-alpha, IFN- gamma, CCL2, and GM-CSF levels (pg/mL) in supernatant collected from NK and SKOV3 coculture were analyzed by Luminex analysis. The mean ± SD from three replicates is shown. Statistical significance is determined by two-way ANOVA with Dunnett’s multiple comparisons test between mock and SKOV3 coculture vs. NK-CAR and SKOV3 coculture. *P* values are indicated on the graph. Data are representative of two independent experiments.

Notably, a similar strategy can be used to design receptors for selective expression on the surface of immune cells such as T cells. A previous study showed that a CAR composed of anti-CD19 scFv fused to T cell receptor (TCR) subunit CD3ε could only be expressed in T cells due to the requirement for interactions with other TCR subunits (27). We also showed that transfection of CD3ε-based CAR mRNA in activated T cells resulted in high surface expression and tumor- specific killing (Supplementary Fig 4A and B). Thus, utilizing endogenous signaling-receptor biology, we can optimize CAR designs that enables engineering of specific cell types in vivo.

Collectively, results presented here demonstrated the feasibility of engineering primary NK cells with CARs based on the NCR family of receptors. These CAR-NK receptors do not contain intracellular signaling motifs but can engage and activate their natural adaptor such as FcRY and DAP12 to initiate tumor killing and cytokine response. In addition, we found that the expression of the NKp44-based CAR strictly requires the presence of the signaling adaptor DAP12, and can not be stably expressed in non-immune cells. This characteristic enables the delivery of mRNA/LNP encoding NKp44 CAR systemically to achieve specific engineering of NK cells in vivo. These results support the further development of CARs for in vivo delivery that harness regulation in receptor stability for cell selective expression. An mRNA/LNP based delivery of NKp44 based CAR for cancer treatment is currently under clinical development.

## ACKNOWLEDGEMENT

We thank members of the discovery research and translational medicine group at Myeloid Therapeutics for their assistance with experiments and helpful discussion.

