## Supplemental Figures 1-4 for "Chimeric Antigen Cytotoxic Receptors for In-Vivo Engineering of Tumor-targeting Natural Killer Cells"

**Supplemental Figure 1**

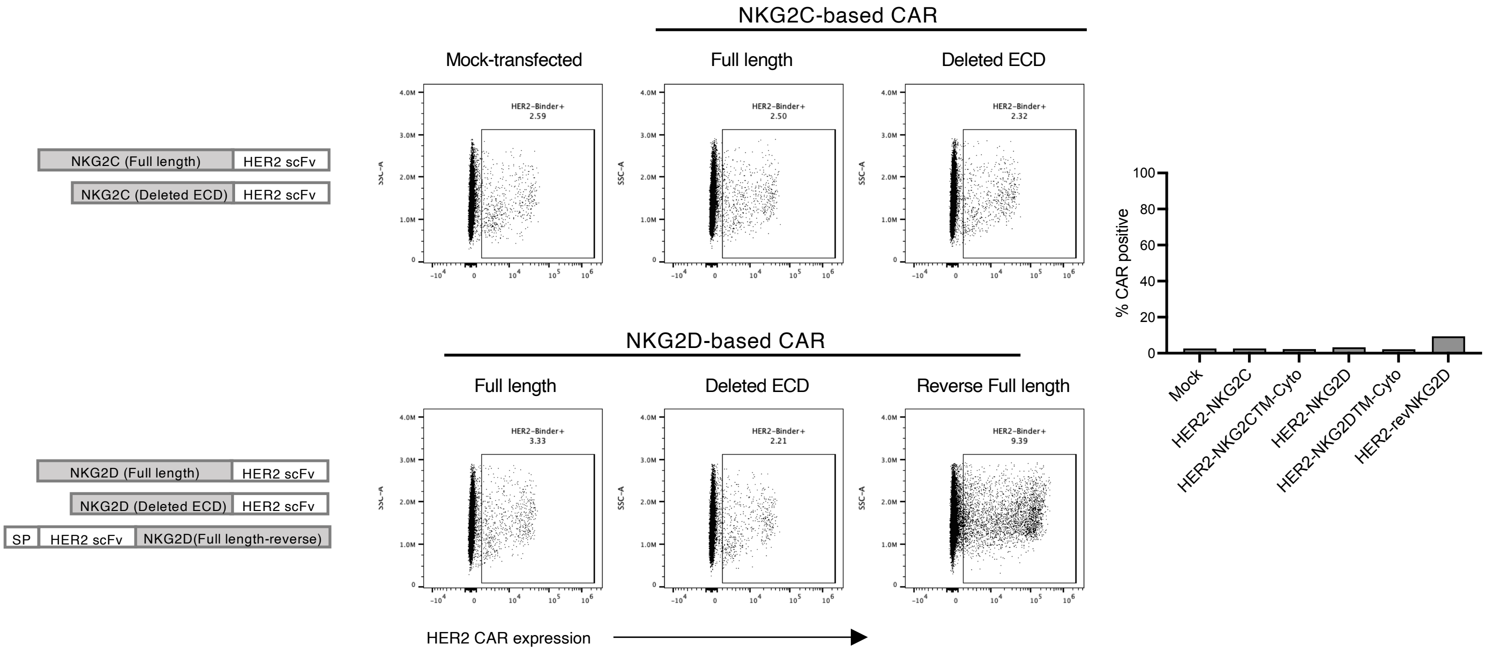


**Supplemental Figure. 1:** Expression of NK-CARs in peripheral blood NK cells. Schematic representation of the designed NK-CARs and cell-surface expression in primary NK cells 20 hours after transfection with NKG2C-based receptors and NKG2D-based receptors. Cell-surface expression of CARs was determined by labeling with recombinant HER2-conjugated with AF647 dye. The graphs show the percentage CAR positive for the different NK-CARs. SP indicates signal peptide.

**Supplemental Figure 2**

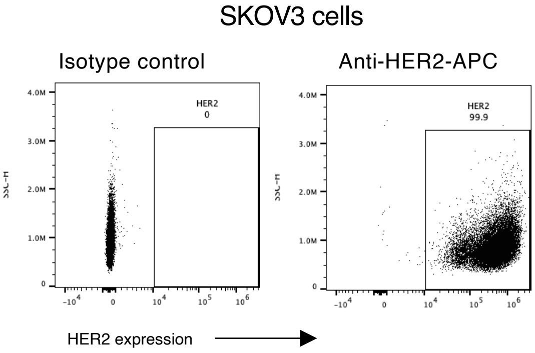


**Supplemental Figure. 2:** Expression of surface antigen HER2 in SKOV3 ovarian cancer cells. Cell-surface expression was determined by labeling with anti-HER2 antibody conjugated with APC dye.

**Supplemental Figure 3**

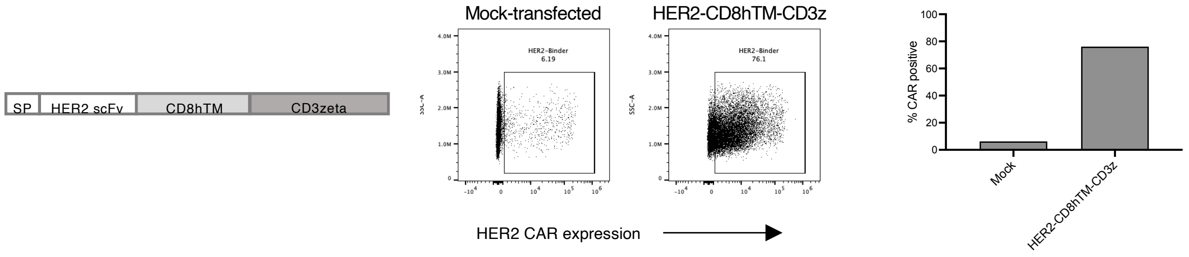


**Supplemental Figure. 3:** Expression of CD3ζ NK-CAR in peripheral blood NK cells. Schematic representation of the designed NK-CAR and cell-surface expression in primary NK cells 20 hours after transfection with CD3ζ-based receptors. Cell-surface expression of CAR was determined by labeling with recombinant HER2-conjugated with AF647 dye. The graph shows the percentage CAR positive. SP indicates signal peptide. Data are representative of three independent experiments in two independent donors.

### Supplemental Figure 4

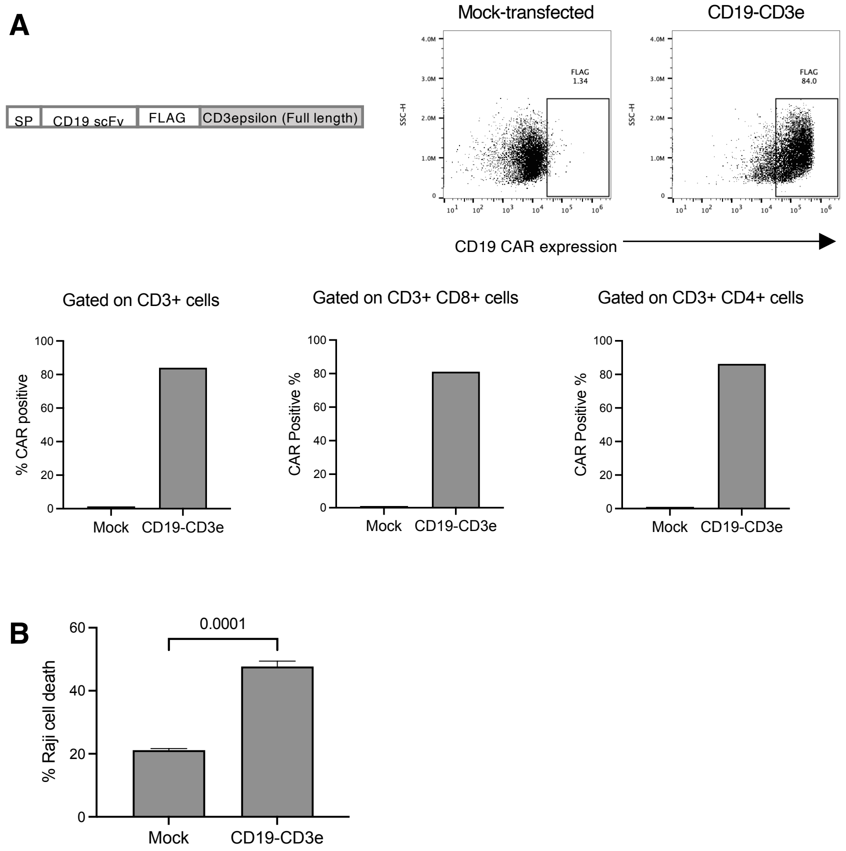


**Supplemental Figure. 4:** Expression and activity of CD19–CD3□ fusion CAR in peripheral blood T cells. (A) Schematic representation of the designed CD19–CD3□ fusion CAR and cell-surface expression in primary T cells 20 hours after transfection. Cell-surface expression of CAR was determined by labeling with anti-FLAG BV421 antibody. The graphs show the percentage CAR-positive population in CD3+, CD3+ CD8+, and CD3+ CD4+ cell populations. SP indicates signal peptide. (B) Peripheral blood T cells transfected with CD19-CD3□ fusion CAR were cocultured with CD19+ Raji tumor cells at a 5:1 E:T ratio for 40 hours at 37□C. The graph shows the percentage Raji-specific cell death. The mean ± SD is plotted, and statistical significance is determined by the unpaired t-test between mock-transfected vs. CD19–CD3□ fusion CAR-transfected T cells. P values are indicated on the graph. Representative data from two independent experiments are shown.
